# Transcriptional landscape in *BRAF* wild type metastatic melanoma

**DOI:** 10.1101/2022.03.31.486532

**Authors:** Elena Lastraioli, Federico Alessandro Ruffinatti, Francesco di Costanzo, Luca Munaron, Annarosa Arcangeli

**Affiliations:** Department of Experimental and Clinical Medicine, University of Florence, Viale GB Morgagni 50, 50134 Florence, Italy; Department of Life Sciences and Systems Biology, University of Torino, Via Accademia Albertina 13, 10123 Torino, Italy; Medical Oncology Unit, Azienda Ospedaliero-Universitaria Careggi, Largo Brambilla 3, 50134 Florence, Italy; Complex Dynamics Study Centre (CSDC), University of Florence

**Author notes:** Equally contributed to the paper (Elena Lastraioli, Federico Alessandro Ruffinatti).

**Keywords:** metastatic melanoma, wild type *BRAF*, transcriptomics, microdissection, ribosomes

## Abstract

Melanoma is a relative rare disease worldwide, nevertheless it has a great relevance in some countries such as in Europe. In order to shed some light upon the transcriptional profile of skin melanoma, we compared the gene expression of 6 independent tumours (all progressed towards metastatic disease and with wild type *BRAF*) to the expression profile of healthy melanocytes. Paraffin-embedded samples were manually microdissected to obtain enriched samples and then RNA was extracted and analysed through a microarray-based approach. An exhaustive bioinformatics analysis was performed to identify differentially expressed transcripts between the two groups as well as enriched functional terms. Overall, 50 up- and 19 down-regulated transcripts were found to be significantly changed in tumour compared to control tissue. Among the upregulated transcripts, the majority belonged to the immune response group and to proteasome, while most of the downregulated genes were related to cytosolic ribosomes. Interestingly, Gene Set Enrichment Analysis (GSEA), along with the analysis of the expression data retrieved from TCGA/GTEx databases, confirmed the general trend of downregulation affecting cytoribosome proteins. In contrast, transcripts coding for mitori-bosome proteins showed the opposite trend.

## 1. Introduction

According to the definition reported by the Dictionary of Cancer Terms (https://www.cancer.gov/publications/dictionaries/cancer-terms) melanoma is “A form of cancer that begins in melanocytes (cells that make the pigment melanin). It may begin in a mole (skin melanoma), but can also begin in other pigmented tissues, such as in the eye or in the intestines”.

Overall, analysing global incidence and mortality worldwide, melanoma is a relatively rare disease that affected 325,000 people in 2020 and 57,000 died because of the disease (source: Globocan 2020, https://gco.iarc.fr/today/online-analysis-table). Nevertheless, skin melanoma has a great relevance in certain countries such as Australia and Europe, where the incidence is higher. In Europe, skin melanoma represents the 7^th^ most frequent malignancy and accounts for roughly 46% of the incident cases in the whole world (https://gco.iarc.fr/today/data/factsheets/cancers/16-Melanoma-of-skin-fact-sheet). Melanocytes can give rise to benign lesions called melanocytic naevi than can progress towards malignant lesions termed melanomas. Melanomas are classified according to TNM staging system (AJCC staging manual 8^th^ edition, issued in 2016 and updated in 2018) [1], and a global melanoma database has been released [2]. Data obtained from clinical and pathological evaluation of melanoma are combined to divide patients into staging groups with different outcomes [3]. In addition, specific classification systems for melanoma have been defined by Clark [4] and Breslow [5] long ago.

The expression profiles and somatic mutations of advanced lesions and metastases have been defined and are reported in the Cancer Genome Atlas Network [6], while less is known about the initial phases of melanoma progression [7]. The most frequent genetic alteration described in melanoma is *BRAF* mutation, present in roughly 50% of the patients, according to the COSMIC database (Catalogue Of Somatic Mutations In Cancer) [8]. The vast majority of *BRAF* mutations is represented by a missense mutation named V600E leading to the substitution of a glutamic acid with a valine in codon 600 [6,9]. The final phenotype is characterized by the constitutive activation of the mitogen-activated protein kinase (MAPK) pathway sustaining cell proliferation and preventing apoptosis. BRAF inhibitors (such as Vemurafenib and Dabrafenib) have been approved for the treatment of metastatic melanoma [10] since they significantly improve progression-free and overall survival although resistance is rapidly acquired [11]. Currently, for patients not carrying *BRAF* mutations no target therapy is available, therefore a significant effort is needed to better define their molecular profile in order to identify potential molecular markers and targets for therapy. Therapy for this group of patients mainly relies on immunotherapy and checkpoint inhibitors [12–14].

The aim of this study was to analyse the transcriptomic profile of patients suffering from metastatic melanoma without *BRAF* mutations in order to evaluate the possible differences in the expression profile between advanced melanoma cells and healthy melanocytes.

## 2. Results

### 2.1 RNA extraction and array hybridisation

In order to obtain a comparative transcriptomic profile of melanoma cells relative to the healthy tissue (i.e., non-dysplastic naevi), we performed manual microdissection of paraffin-embedded surgical samples [15] of both normal tissue and *BRAF* wild type melanoma from patients whose clinicopathological characteristics are in Table 4 (Materials and Methods). Samples hence enriched in the melanocytic population (Figure 1) were processed for RNA extraction and then hybridized on Agilent arrays after RNA validation (see details in Materials and Methods).

**Figure 1.**
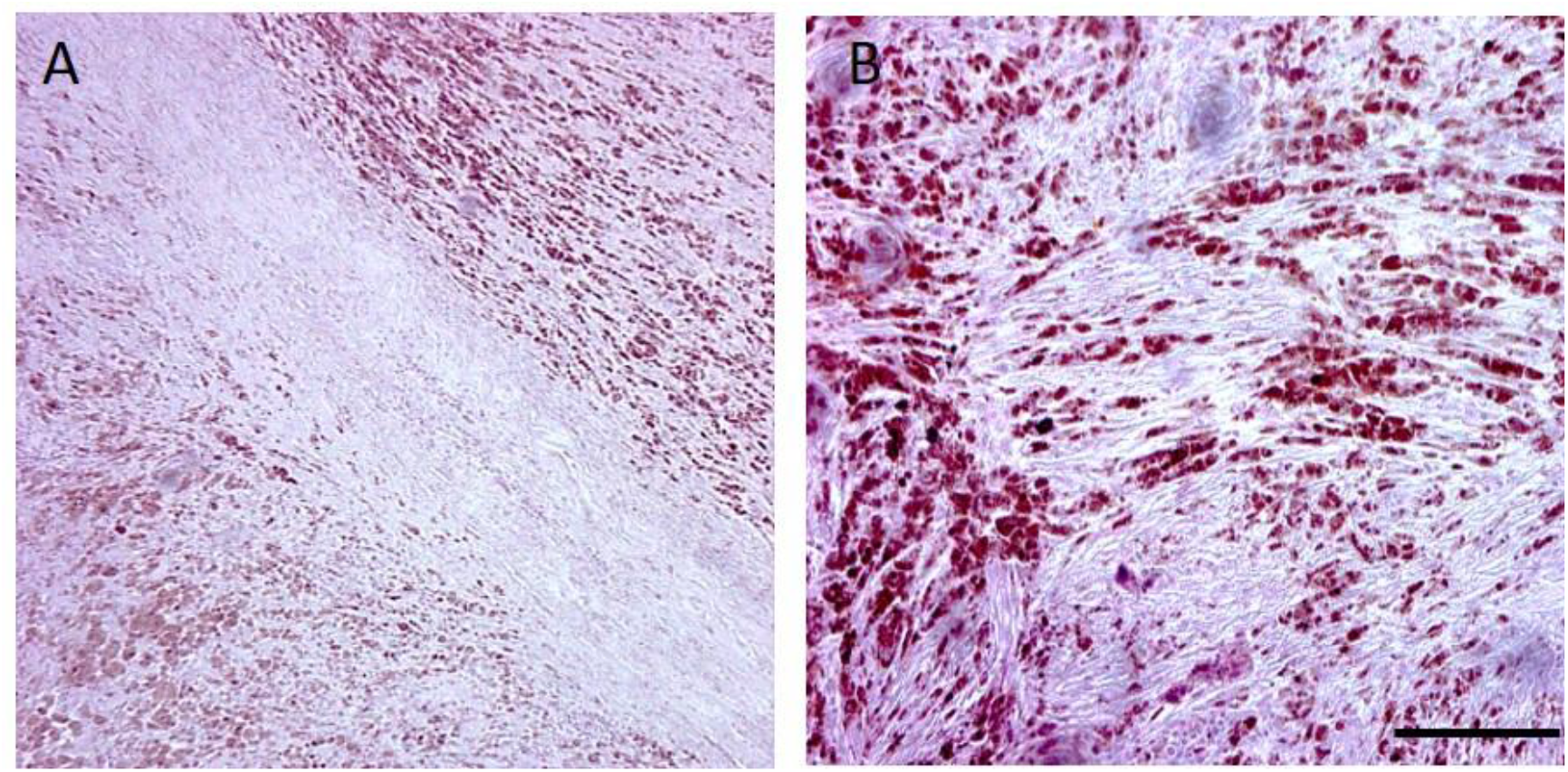
Histopathological microphotograph of a representative melanoma sample. Haematoxylin-Eosin staining of a representative melanoma sample showing areas rich of melanin-producing tumour cells (brown coloured). Scale bar: 50 µm.

### 2.2 Differential Expression Analysis

Gene expression data from microarray experiments were preprocessed according to a standard pipeline, as described in Materials and Methods. The so obtained normalized log_2_ expression data were then filtered and subjected to differential expression analysis (DEA) using the rank-product statistics (see Materials and Methods for details). In particular, *n* = 6 independent samples of *BRAF* wild type metastatic melanoma were compared to a reference array representing a pool of healthy tissues enriched with normal melanocytes (biological average of *n* = 4 independent samples). Transcripts featuring a BH-FDR *q*-value < 0.05 and a |log_2_FC|> 0.5 were deemed as differentially expressed genes (DEGs). Overall, DEA returned a list of 84 statistically significant probes, but just 69 of them could be annotated. Specifically, 50 up- and 19 down-regulated transcripts were found to be significantly changed in tumour compared to control tissue, as reported in Table 1 and Table 2 respectively.

**Table 1.** Upregulated genes, as resulted from the statistical comparison of melanoma *vs* healthy samples (Rank Product). Positive log_2_FC values indicate overexpression in tumour compared to healthy.

| Probe ID | GeneSymbol | Description | Log2FC | q-value<br>BH-FDR | P.value |
| --- | --- | --- | --- | --- | --- |
| A_32_P137939 | ACTB | actin beta | 1.294547234 | 9.182E-08 | 5.459E-11 |
| A_33_P3223592 | APOE | apolipoprotein E | 1.902652471 | 2.359E-09 | 3.507E-13 |
| A_33_P3378531 | AS3MT | arsenite methyltransferase | 1.071076162 | 0.00000902 | 1.073E-08 |
| A_33_P3296198 | C5orf63 | chromosome 5 open reading frame 63 | 1.37517848 | 0.0000034 | 2.527E-09 |
| A_33_P3292854 | CALR | calreticulin | 0.641546029 | 0.0004772 | 0.000001738 |
| A_33_P3280066 | CAVIN1 | caveolae associated protein 1 | 0.917622702 | 0.00002915 | 5.633E-08 |
| A_33_P3284508 | CD14 | CD14 molecule | 1.828631582 | 4.271E-09 | 1.27E-12 |
| A_33_P3229196 | CD151 | CD151 molecule (Raph blood group) | 0.633198686 | 0.0005929 | 0.000002379 |
| A_33_P3252612 | CYP2W1 | cytochrome P450 family 2 subfamily W member 1 | 0.545968369 | 0.001773 | 0.000009091 |
| A_24_P100673 | EMC4 | ER membrane protein complex subunit 4 | 0.676267408 | 0.004377 | 0.0000283 |
| A_33_P3333455 | EMILIN1 | elastin microfibril interfacier 1 | 0.535057595 | 0.0003992 | 0.000001306 |
| A_33_P3379436 | FAM74A4 | family with sequence similarity 74 member A4 | 0.847195187 | 0.00003356 | 6.985E-08 |
| A_32_P342064 | FTH1 | ferritin heavy chain 1 | 0.504874589 | 0.005501 | 0.00003802 |
| A_23_P13899 | GAPDH | glyceraldehyde-3-phosphate dehydrogenase | 0.612588407 | 0.0003961 | 0.000001325 |
| A_33_P3585268 | GNAI2 | G protein subunit alpha i2 | 1.331705216 | 1.009E-07 | 5.25E-11 |
| A_24_P108451 | GPI | glucose-6-phosphate isomerase | 0.834306703 | 0.000005515 | 4.509E-09 |
| A_33_P3354322 | GPX1 | glutathione peroxidase 1 | 0.840765337 | 0.00004531 | 1.044E-07 |
| A_33_P3287218 | GSTK1 | glutathione S-transferase kappa 1 | 0.587845009 | 0.0002005 | 5.364E-07 |
| A_33_P3379962 | HLA-A | major histocompatibility complex, class I, A | 0.863145543 | 0.001296 | 0.000006167 |
| A_33_P3424803 | HLA-C | major histocompatibility complex, class I, C | 0.763836459 | 0.001062 | 0.000004734 |
| A_23_P162874 | HSP90AA1 | heat shock protein 90 alpha family class A member 1 | 0.616357697 | 0.0003777 | 0.000001207 |
| A_23_P72737 | IFITM1 | interferon induced transmembrane protein 1 | 0.806530289 | 0.000005975 | 5.773E-09 |
| A_24_P605563 | IGLC1 | immunoglobulin lambda constant 1 | 0.517023672 | 0.0005772 | 0.000002231 |
| A_23_P167168 | JCHAIN | joining chain of multimeric IgA and IgM | 0.5290713 | 0.002256 | 0.00001224 |
| A_32_P452655 | LGALS9C | galectin 9C | 0.605020153 | 0.0001375 | 3.578E-07 |
| A_23_P91619 | MIF | macrophage migration inhibitory factor | 0.956800118 | 0.00000965 | 1.219E-08 |
| A_23_P1904 | MS4A2 | membrane spanning 4-domains A2 | 0.630405471 | 0.00002615 | 4.471E-08 |
| A_23_P106844 | MT2A | metallothionein 2A | 0.551885103 | 0.0004391 | 0.000001534 |
| A_33_P3239879 | NAA38 | N-alpha-acetyltransferase 38, NatC auxiliary subunit | 0.782408565 | 0.00004663 | 0.000000104 |
| A_23_P33022 | POLR2L | RNA polymerase II, I and III subunit L | 0.83673208 | 0.00001238 | 1.749E-08 |
| A_33_P3377199 | PRDX1 | peroxiredoxin 1 | 0.72334012 | 0.0003411 | 0.000001065 |
| A_33_P3234899 | PSMB3 | proteasome 20S subunit beta 3 | 0.794103899 | 0.00009369 | 2.298E-07 |
| A_23_P65427 | PSME2 | proteasome activator subunit 2 | 0.701327257 | 0.00001028 | 1.375E-08 |
| A_23_P434301 | PTMA | prothymosin alpha | 0.582074221 | 0.0004559 | 0.000001626 |
| A_33_P3382595 | RN7SK | RNA component of 7SK nuclear ribonucleoprotein | 0.679874148 | 0.00002798 | 4.367E-08 |
| A_23_P69431 | RPL4 | ribosomal protein L4 | 0.508935662 | 0.0009252 | 0.000004057 |
| A_23_P106708 | RPS2 | ribosomal protein S2 | 0.663572335 | 0.004081 | 0.00002457 |
| A_23_P372874 | S100A13 | S100 calcium binding protein A13 | 1.043344505 | 0.00002044 | 3.038E-08 |
| A_24_P261169 | SEMA4D | semaphorin 4D | 0.574343754 | 0.0000385 | 8.299E-08 |
| A_33_P3413989 | SERPING1 | serpin family G member 1 | 1.286289758 | 6.052E-07 | 4.048E-10 |
| A_23_P95213 | SFTPC | surfactant protein C | 0.854394678 | 0.005545 | 0.00003874 |
| A_33_P3481987 | SLC16A12 | solute carrier family 16 member 12 | 0.850399397 | 0.00003454 | 6.93E-08 |
| A_33_P3388491 | SLC66A1 | solute carrier family 66 member 1 | 1.439889048 | 0.00000662 | 7.381E-09 |
| A_33_P3587376 | SNAR-A3 | small NF90 (ILF3) associated RNA A3 | 1.423309237 | 0.0000895 | 2.129E-07 |
| A_33_P3370461 | SUZ12P1 | SUZ12 pseudogene 1 | 0.916094423 | 0.00003032 | 5.633E-08 |
| A_33_P3332690 | SUZ12P1 | SUZ12 pseudogene 1 | 0.597411988 | 0.0002652 | 7.688E-07 |
| A_33_P3274199 | TP53I13 | tumor protein p53 inducible protein 13 | 0.579180148 | 0.0004745 | 0.000001763 |
| A_23_P325654 | TRIM42 | tripartite motif containing 42 | 0.925239905 | 0.04699 | 0.0009395 |
| A_33_P3409062 | TYROBP | transmembrane immune signaling adaptor TYROBP | 1.451540833 | 6.771E-08 | 2.516E-11 |
| A_24_P101391 | YBX1 | Y-box binding protein 1 | 0.713695815 | 0.0002987 | 9.101E-07 |

**Table 2.** Downregulated genes, as resulted from the statistical comparison of melanoma *vs* healthy samples (Rank Product). Positive log_2_FC values indicate overexpression in tumour compared to healthy.

| Probe ID | GeneSymbol | Description | Log2FC | q-value<br>BH-FDR | P.value |
| --- | --- | --- | --- | --- | --- |
| A_23_P114445 | MAGEE1 | MAGE family member E1 | -0.507993024 | 0.00005469 | 1.301E-07 |
| A_23_P112774 | PTP4A3 | protein tyrosine phosphatase 4A3 | -0.512513651 | 0.00002708 | 4.227E-08 |
| A_33_P3332348 | RN7SL1 | RNA component of signal recognition particle 7SL1 | -0.802637816 | 0.000002856 | 4.245E-10 |
| A_33_P3244165 | RNA28SN5 | RNA, 28S ribosomal N5 | -1.378194265 | 0.000006921 | 2.058E-09 |
| A_33_P3346552 | RNA28SN5 | RNA, 28S ribosomal N5 | -1.012460836 | 0.0000205 | 2.894E-08 |
| A_33_P3279708 | RNU2-2P | RNA, U2 small nuclear 2, pseudogene | -0.953718497 | 7.281E-07 | 5.411E-11 |
| A_23_P217068 | RPL12 | ribosomal protein L12 | -0.752923168 | 0.0005958 | 0.000003321 |
| A_24_P142228 | RPL13 | ribosomal protein L13 | -0.578413127 | 0.00000201 | 4.482E-10 |
| A_32_P184518 | RPL21 | ribosomal protein L21 | -0.807427059 | 0.00000608 | 3.615E-09 |
| A_32_P118258 | RPL21 | ribosomal protein L21 | -0.861447625 | 0.00002754 | 4.093E-08 |
| A_24_P213783 | RPL31 | ribosomal protein L31 | -0.94369394 | 0.000006654 | 4.451E-09 |
| A_23_P18142 | RPL32 | ribosomal protein L32 | -0.567895777 | 0.001532 | 0.00001196 |
| A_33_P3329916 | RPL6 | ribosomal protein L6 | -0.803141199 | 0.000007194 | 3.208E-09 |
| A_32_P857658 | RPLP1 | ribosomal protein lateral stalk subunit P1 | -0.735043282 | 0.0003437 | 0.000001481 |
| A_23_P147888 | RPLP2 | ribosomal protein lateral stalk subunit P2 | -0.511690664 | 0.0009097 | 0.000006017 |
| A_24_P418418 | RPS17 | ribosomal protein S17 | -0.930710287 | 0.0000174 | 2.07E-08 |
| A_23_P116694 | RPS26 | ribosomal protein S26 | -0.539100203 | 0.00001758 | 1.96E-08 |
| A_33_P3221680 | RPS28 | ribosomal protein S28 | -0.750493979 | 0.0000148 | 1.43E-08 |
| A_23_P46182 | RPS8 | ribosomal protein S8 | -0.757306042 | 0.0000937 | 2.855E-07 |

### 2.3 Genes related to antigen processing and presentation are upregulated in tumor vs control

DEGs that were found to be upregulated in metastatic melanoma samples compared to non-dysplastic controls were tested for functional enrichment using ToppFun web tool (https://toppgene.cchmc.org/, accessed on 22 February 2022). The full table of the statistically significant terms retrieved from such a query can be found as Table SM1 in Supplementary Material section. Inspecting the results, it is noticeable how the top-most ranked functional terms and pathways were almost all related to the positive regulation of some features of the immune response, involving 32 out of the 50 up-regulated DEGs resulting from DEA. For example, the most relevant GO-terms referring to biological processes (BPs) were *innate immune response, defence response to other organism, regulation of immune system process, cell activation, response to external biotic stimulus, leukocyte mediated immunity, antigen processing and presentation of exogenous peptide antigen via MHC class I* (BH-FDR < 1.7 · 10^−4^). Accordingly, the most significant KEGG pathway was *antigen processing and presentation* (BH-FDR = 6.4 · 10^−4^), accounting for 5 DEGs having a central role in MHC class I pathway: *HLA-A, HLA-C, CALR, PSME2, HSP90AA1* (Figure 2, genes in magenta).

**Figure 2.**
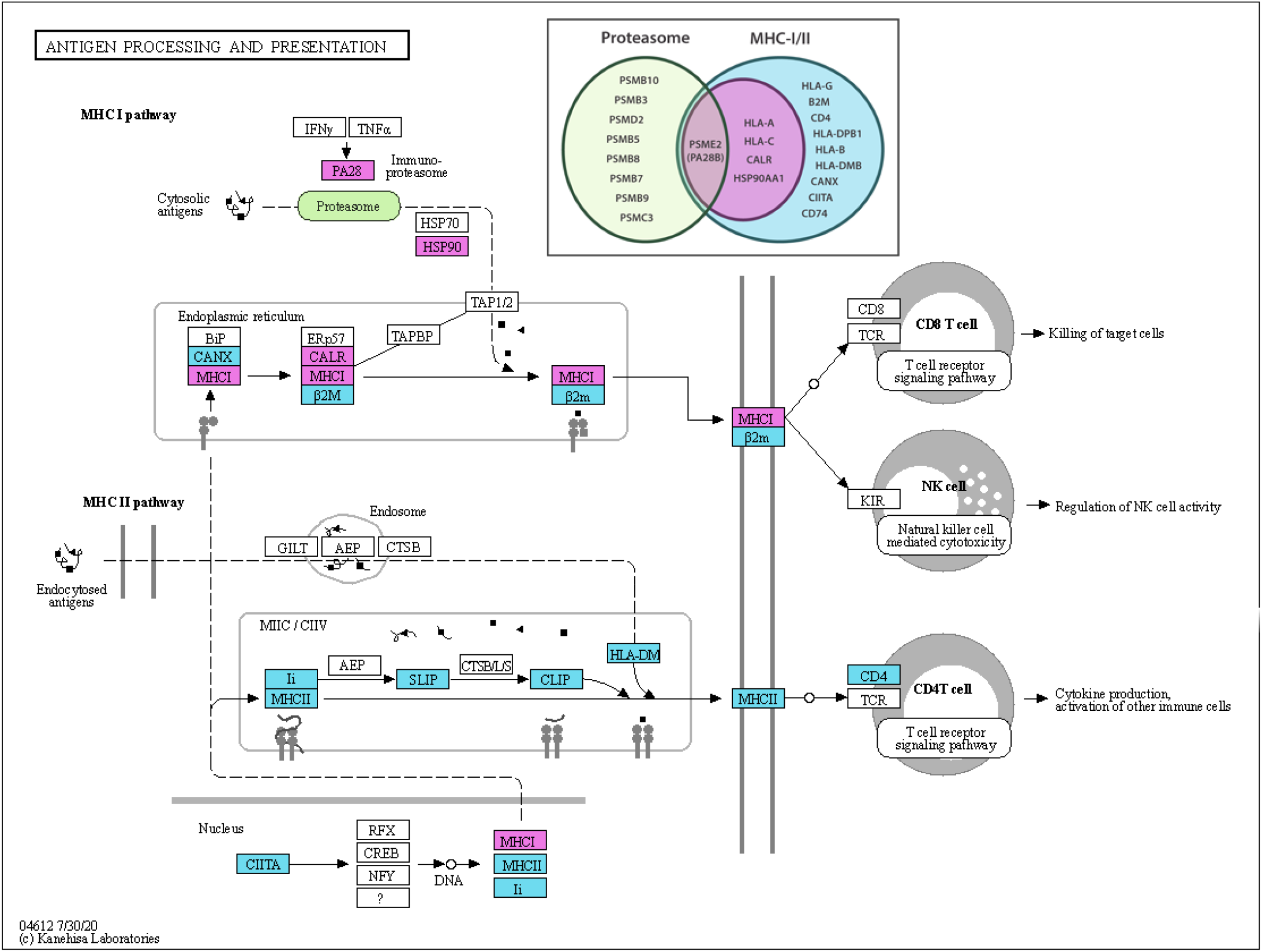
Illustration of the antigen processing and presentation KEGG pathway. Upregulated DEGs detected by rank product statistics are filled with magenta (*HLA-A* and *HLA-C* are here collectively referred to as *MHCI*; *HPS90* is a short for *HSP90AA1*; *PA28* is an alias for *PSME1-2-3*). In cyan are the elements of the pathway additionally detected by GSEA leading-edge analysis. GSEA also revealed a significant involvement of the proteasome complex (in green). Using the same colour-code, the Venn diagram in the upper inset shows the complete lists of the official gene symbols found to be upregulated within the two KEGG pathways.

Such a finding was confirmed by GSEA (see Materials and Methods), according to which the gene set corresponding to that pathway was positively enriched (NES = 2.03, FDR *q*-value = 0.064, Figure 3A). More in detail, the leading-edge analysis identified 13 main genes involved in the upregulation of both the MHC class I and class II pathways, thus extending the previous set of 5 DEGs detected on the basis of a gene-wise hypothesis testing (Figure 2, genes in magenta plus genes in cyan). In addition, GSEA pointed at a significant positive regulation of the *proteasome* complex (NES = 2.07, FDR *q*-value = 0.069, Figure 3B), another KEGG-pathway term deeply connected to the previous one, with a leading-edge featuring 9 genes coding for different proteasome subunits, including the pivotal proteasome activator subunit 2 (*PSME2*) as a linker between the two gene sets (Figure 2, green ellipse and Venn diagram).

**Figure 3.**
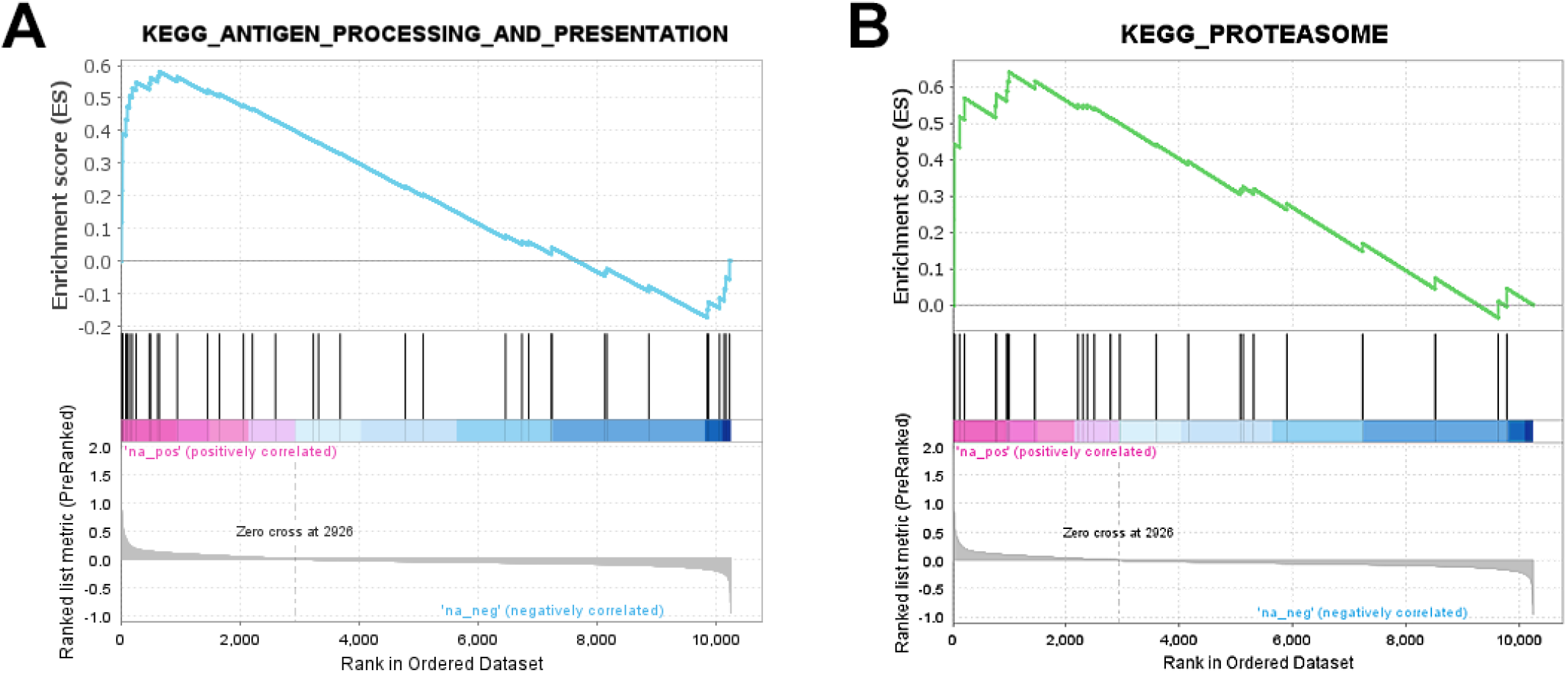
GSEA enrichment plots. Profile of the running ES score (upper boxes) and positions of gene set members on the rank-ordered list from microarray experiments (lower boxes) for **(A)** *antigen processing and presentation* and **(B)** *proteasome* KEGG pathways, respectively. The leading-edge comprises that portion of the gene set between the (absolute) ES maximum and the nearest edge of the ranked list.

### 2.4 Cytosolic ribosome proteins are downregulated in tumor vs control

Strikingly, the vast majority of the transcripts found to be downregulated in metastatic melanoma compared to the control reference were related to cytosolic ribosomes (see Table 2). More in detail, 13 out of the 19 downregulated DEGs corresponded to ribosomal proteins (rProteins) of either the large (60S) or the small (40S) cytosolic ribosome subunit. In addition, two different probes targeting the product of the RNA28SN5 gene—the ribosomal RNA giving rise to the 28S subunit—were among the 19 DEGs featured by the list of downregulated genes.

In order to confirm and extend these results, we ran a GSEA testing the whole spectrum of rProteins, of both cytosolic and mitochondrial origin. To do this, we took advantage of the already available Ribosomal Protein Gene Set (RPGS), that is the complete list of all human gene symbols related to ribosomes we assembled for a recent work in order to answer a similar scientific question [15]. Such an analysis confirmed a significant downregulation of the structural constituents of both the large (60S) and the small (40S) cytosolic ribosome subunits (Figure 4A-C). On the contrary, and most interestingly, mitochondrial rProteins and rRNA did not show any significant downregulation, but rather an opposite trend (Figure 4D-F).

**Figure 4.**
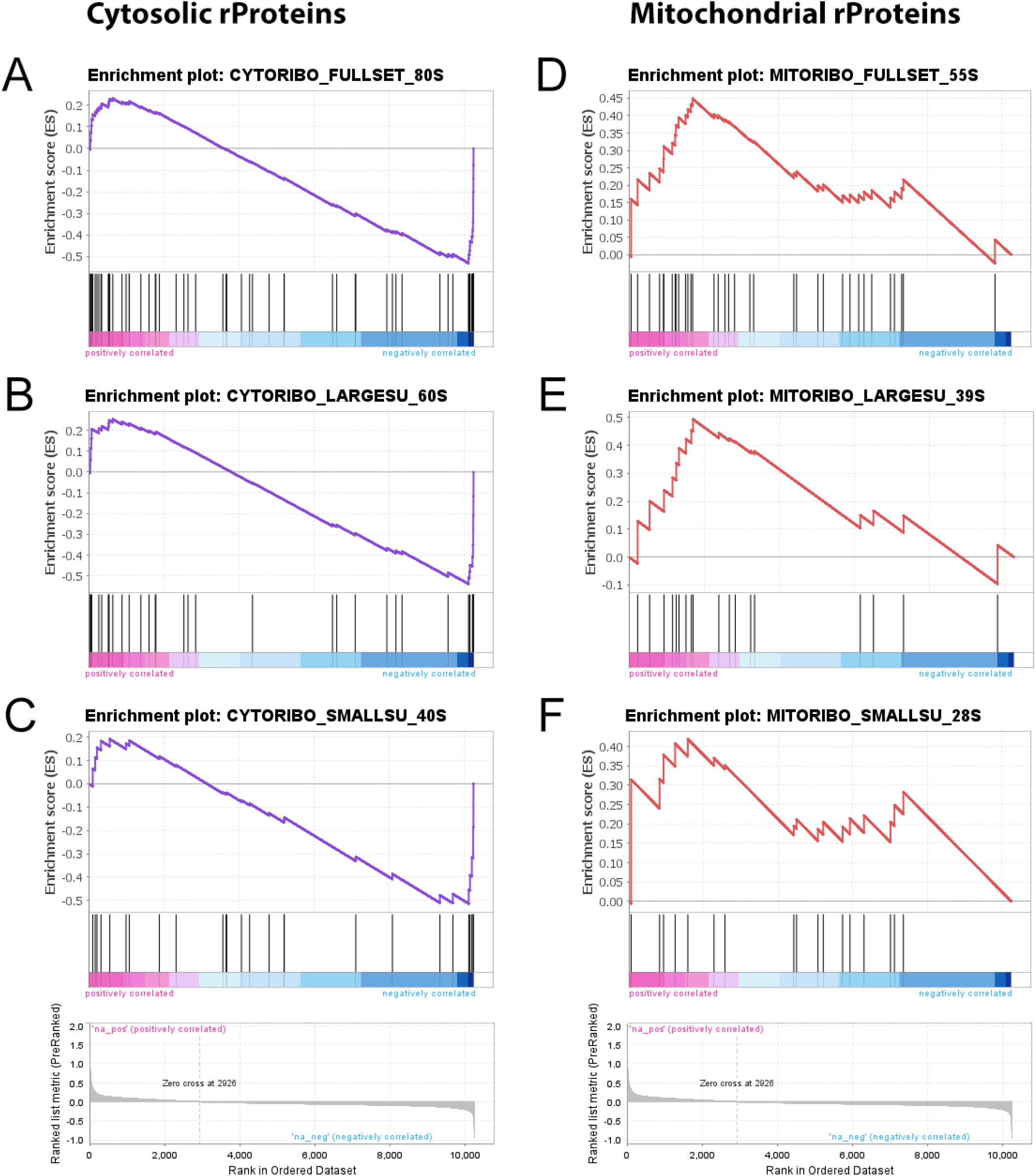
Gene set enrichment analysis of the ribosomal protein gene set. **(A–C)** Downregulated cytosolic rProtein transcripts were significantly enriched (*q*-values: 1.5 · 10^−4^, 0.002, 0.013 for 80S, 60S, and 40S subunit gene sets, respectively). **(D, E)** In contrast, mitochondrial rProtein genes showed a consistent upregulation (*q*-values: 0.205, 0.151, 0.231 for 55S, 39S, and 28S subunit gene sets, respectively).

### 2.5 External validation through TCGA vs GTEx cohorts

In order to rule out any technical artifact related to microarray hybridization or sample origin, we decided to externally validate these findings using UCSC Xena Browser (University of California, Santa Cruz, http://xena.ucsc.edu/, accessed on 24 February 2022) that provides a convenient way to access gene expression data stored in TCGA database for the comparative analysis of tumour samples with the normal analogous available from GTEx database (https://gtexportal.org/home/) [16,17]. TCGA samples were thus filtered based on cancer type (Skin Cutaneous Melanoma, SKCM), stage (metastatic), and genomic subtype (*BRAF* wild type). The so obtained cohort featured 179 SKCM samples that were compared with the corresponding healthy GTEx cohort of skin normal tissue made of 557 samples, for a total sample size of *n* = 736.

A thorough validation was carried out for the following sets of genes resulting from the corresponding GSEA leading-edge analysis showed in Figures 3-4: *MHC pathway* (14 genes), *proteasome* (9 genes), *cytosolic rProteins* (14 genes), and *mitochondrial rProteins* (14 genes). Notably, almost all gene differential expression we tested could be confirmed by TCGA/GTEx data in terms of both change direction (log_2_FC sign) and statistical significance (Figure 5). The detailed validation scores are given in Table 3.

**Table 3.** Number of genes subjected to validation by TCGA/GTEx databases and their outcomes.

| Pathway Name | GSEA<br>leading-edge | Xena<br>opposite FC | Xena<br>not significant | Xena<br>concordant | Validation score |
| --- | --- | --- | --- | --- | --- |
| MHC pathway | 14 | 1 | 0 | 13 | 92.9 % |
| proteasome | 9 | 0 | 0 | 9 | 100 % |
| cytosolic rProteins | 14 | 1 | 2 | 11 | 78.6 % |
| mitochondrial rProteins | 14 | 0 | 1 | 13 | 92.9 % |

**Figure 5.**
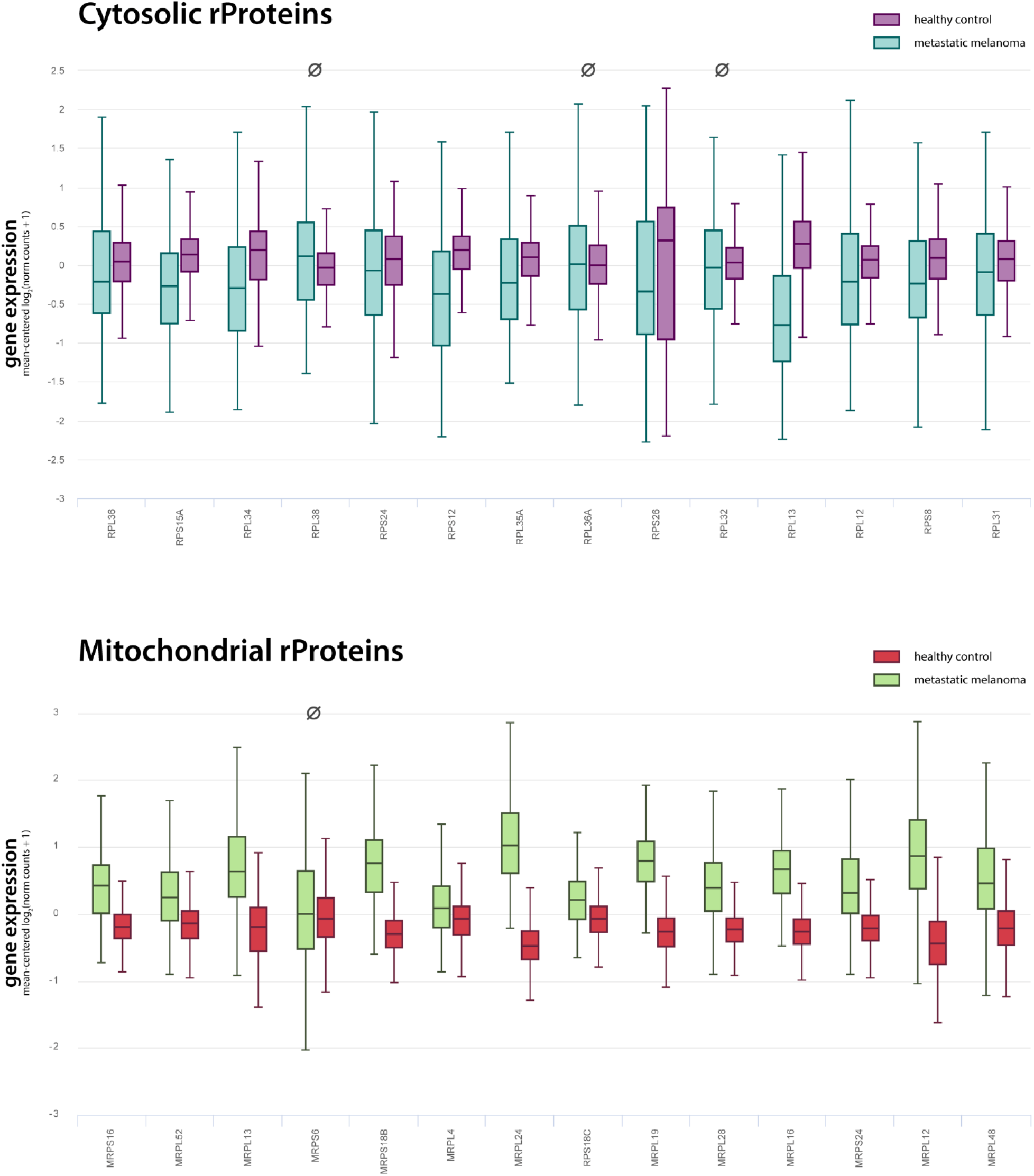
Ribosomal protein expression from TCGA/GTEx databases for metastatic melanoma samples. Xena browser was used to filter TCGA samples and keep only data from Skin Cutaneous Melanoma studies, of metastatic type, and with no mutations on *BRAF* gene. The final cohort featured 179 tumour samples from TCGA and 557 healthy samples from GTEx, for a total sample size of *n* = 736. RNA-Seq expression data are given in units of log_2_ RSEM normalized counts, gene-wise mean-centred, and shown as boxplots for the 14 cytosolic (magenta and teal) and the 14 mitochondrial (red and green) rProteins emerged from the leading-edge analysis of microarray data. Overall, 24 out of 28 differential expressions were confirmed by TCGA data, supporting the evidence of a generalized downregulation of cytosolic rProteins and an overexpression of mitochondrial ones. The four unconfirmed comparisons are marked with the symbol Ø (see Table 1 for more details).

## 3. Discussion

In this study RNA extracted from paraffin-embedded samples of metastatic melanoma was hybridised on Agilent microarrays to evaluate gene expression levels in transformed melanocytes compared to normal tissue.

GSEA and ToppFun functional enrichment analysis of upregulated DEGs showed a consistent involvement of the immune system, with 70% of the overexpressed genes coherently annotated to some immune response-related process. This is in line with the well-known immunogenic nature of melanoma and the recent literature pointing at MHC-I/II protein expression as a powerful prognostic marker to predict the effectiveness of anti-CTLA-4 and anti-PD-1 immunotherapy in metastatic melanoma and other cancer types [18–22].

Even though most of these papers agree on the fact that a transcriptional downregulation of MHC-I and MHC-II genes is a common feature of advanced untreated melanomas, in our study the opposite seems to be true. Notably this cannot be ascribed to some spurious effects induced by drugs—such as antibodies targeting the immune checkpoints—since all the samples we used for RNA extraction were excised from patients before any therapeutic schedule. Moreover, RNA-Seq data from TCGA/GTEx database confirmed such a significant overexpression of all the MHC class I genes (*HLA-A, HLA-B, HLA-C*), as well as MHC class II (*HLA-DMA, HLA-DOA, HLA-DPA1, HLA-DQA1, HLA-DRA*), in *BRAF*-wild type metastatic melanoma compared to healthy skin tissue (Supplementary Figure SM1). Rather, fold-change signs could be dependent on the particular stage at which melanoma samples were collected. Indeed, MHC gene expression profile has already been reported to be heavily dependent on tumour progression and its gradual loss is likely to facilitate the evasion of cancer cells from immune surveillance [23].

The other gene set we found to be upregulated in our cohort of metastatic melanoma patients compared to healthy controls was related to proteasomal function. Beyond its increased expression, proteasome complex in melanoma cells may be also overactive because of the overexpressed *PSME2* gene, the proteasome activator complex subunit 2 (aka *PA28B*), thus contributing in turn to the increased antigen presentation by the MHC class I pathway discussed above (see Figure 2). These data agree with the notion that melanoma cells heavily rely on proteasomal function to survive, so that selective proteasome inhibitors have already been used as new attractive therapeutics for this type of cancer [24– 26].

On the other hand, DEGs we found to be downregulated in melanoma samples compared to healthy controls were mostly related to cytosolic ribosomal proteins (rProteins). This was not completely unexpected given the accumulating evidence that relates cancer onset and progression with alterations of cell translational machinery (see, for instance, [27] for a good review focusing on this fascinating topic). Specifically, both enhanced and reduced ribosome biogenesis and protein synthesis have been reported to be associated with cancer in mammals, depending on the particular type of tissue and stage taken into consideration [28–32]. For this reason, we decided to examine rProteins configuration in metastatic melanoma more in depth by running a GSEA of all the structural constituents of both cytosolic and mitochondrial ribosomes. Intriguingly, the two ribosome types showed opposite patterns of deregulation: while cytosolic rProteins tended to be under expressed in metastatic melanoma, mitochondrial ones were sharply upregulated.

Such a finding is of great interest considering in particular the data we have recently published in another paper addressing transcriptional alterations in colorectal carcinoma [15]. As in the present case, even in that study we were able to find a consistent change in rProtein expression, but with a different FC sign, in that the upregulation of rProteins concerned cytosolic ribosomes and not the mitochondrial ones. Importantly, in both studies, all the ribosomal transcriptional alterations we reported found confirmation in TCGA/GTEx large cohorts of patients.

In the early phases of melanoma pathogenesis, the key metabolic process is represented by glycolysis and after the occurrence of *BRAF* mutations the stimulation of transcription factors acting as key regulators of such process make it even more effective [33,34]. Moreover, in *BRAF* mutated cells the Oxidative Phosphorylation (OXPHOS) is inhibited [35]. It is well-known that between these two metabolic phenotypes a dynamic switch occurs and plasticity plays a key role in melanoma [36–38] leading to metabolic reprogramming of the cells. To make the picture more complex, it has been shown that some melanomas are able to exploit diverse nutrients and energy sources to adapt to different extracellular conditions thus showing a “hybrid” glycolysis/OXPHOS metabolic phenotype [39–41]. Finally, the so-called “Reverse Warburg” effect has been described in melanoma cells [38,40,42]. This effect relies on the stimulation of cancer-associated fibroblasts (CAFs) that increase their glucose upload and lactate secretion through Monocarboxylate Transporter (MCT) family proteins [43]. Moreover, lactate can be internalised by cancer cells *via* MCT and conveyed into the Krebs cycle thus fuelling OXPHOS. In this view, immune cells can deregulate metabolic pathways representing a link between the deregulated pathways emerged in this paper. As a further confirmation the transcript of SLC66A1, encoding an MCT, resulted to be upregulated in our cohort.

All the proteins encoded by mitochondrial DNA are involved in the assemble and functioning of the respiratory complexes along with protein encoded by nuclear DNA. For this reason, the OXPHOS biogenesis is subjected to a synchronized regulation of the mitochondrial and cytoplasmic ribosomes. The interplay between nuclear and mitochondrial components for ribosome production and the consequent synthesis of the various proteins involved in glycolysis and OXPHOS is extremely complex. Since our data derive from a transcriptomic approach, they can’t give insights on the expression and function of the glycolytic and OXPHOS enzymes, therefore no robust hypothesis on functional significance can be proposed. Nevertheless, these data could pave the road for further evaluations in which a proteomic and metabolomic approach can be used (to define the expression levels of key glycolytic/ OXPHOS enzymes) as well as a biochemical and physiological approach.

The results reported in this paper might be relevant for two main reasons: (i) the central role of protein synthesis and energy metabolism in cancer, and (ii) the fact that, despite the many recent reports about cytosolic ribosome aberrant function in cancer, there are still few data about the 55S mitochondrial counterparts and their functional interplay with 80S ribosomes. For example, the evidence of possible mitoribosome onco-patterns could provide a new rationale for the design (or repurposing) of novel antibiotics specific for cancer treatment, a still debated clinical practice [44].

Taken together, our data point at a complete and deep remodelling of protein synthesis and degradation in metastatic melanoma that suggests, alongside biopsy genotypisation, a more integrated evaluation of specific gene expression patterns—in particular those related to MHC, proteasomes, and rProteins—as a practice that could help choosing the most effective treatment in a context of personalised medicine.

## 4. Materials and MethodsPatients

6 patients (2 females, 4 males with mean age at diagnosis 60.3 years, range 46-70) suffering from metastatic melanoma not harbouring *BRAF* mutations were enrolled for the study between April 2016 and October 2018 within the OMITERC study coordinated by Medical Oncology Unit, Azienda Ospedaliero-Universitaria Careggi (Florence). All the patients provided an informed written consent, and the study was approved by the local Ethical Committee of Azienda Ospedaliero-Universitaria Careggi (BIO.16.028 released on October 5^th^ 2016). Paraffin-embedded samples of the primary tumours were retrieved from the archives of the Department of Medical Biotechnologies, University of Siena, Italy. The clinical and pathological features of the patients were defined by experienced medical oncologists and pathologists according to the relevant guidelines (Table 4). Moreover, 4 non dysplastic naevi were also collected from the same institution as above.

**Table 4.**
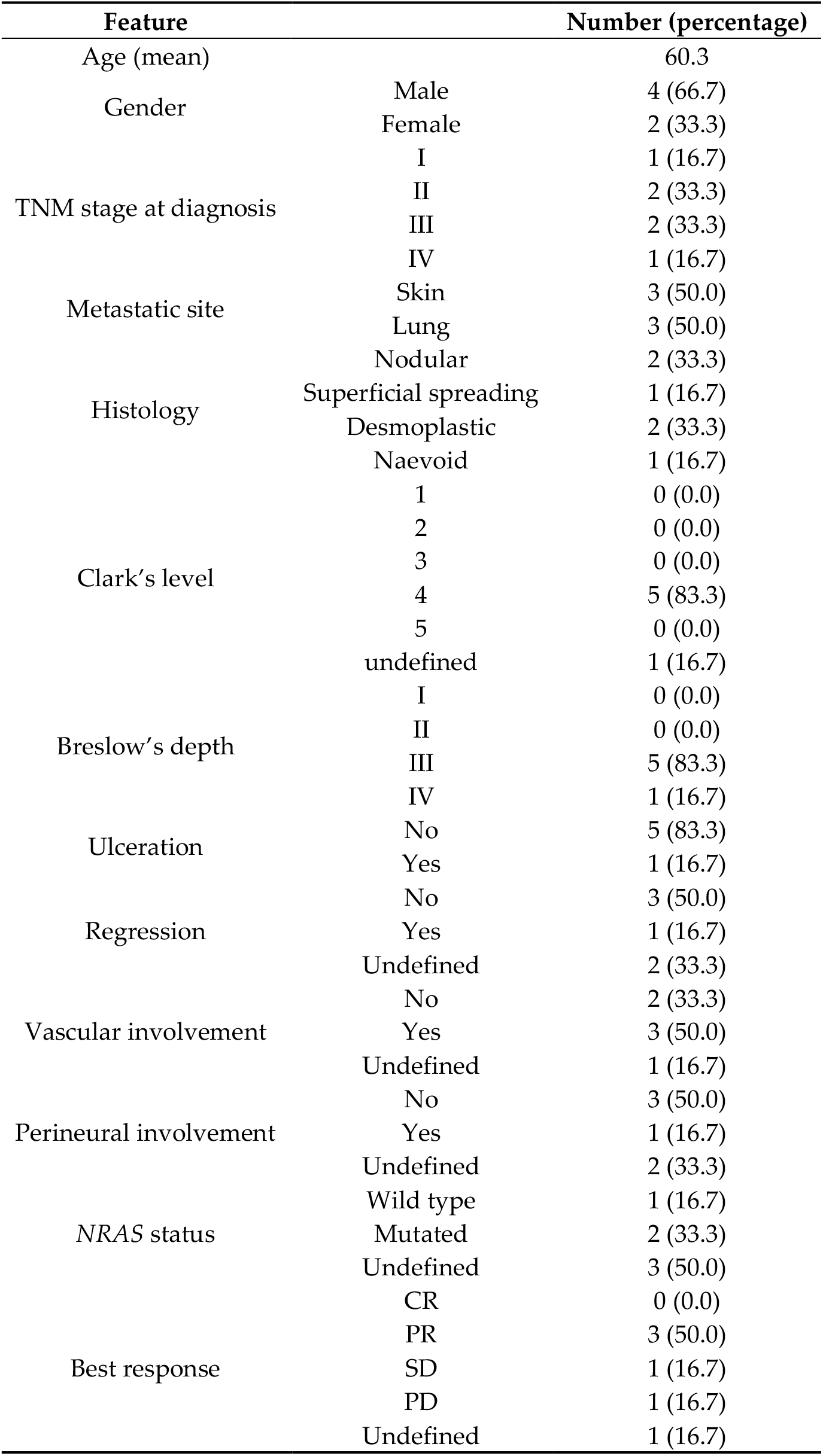
Demographic and clinical features of the patients enrolled in the study.

### Sample preparation

In order to obtain tumour-enriched samples, paraffin-embedded specimens were manually micro-dissected, applying the same protocol as in [15].

### RNA extraction and quality control

Total RNA was extracted from the enriched samples with AllPrep® DNA / RNA FFPE kit (Qiagen) according to the manufacturer’s protocol. The extracted RNA was then checked for its quality and integrity by Agilent 2100 Bioanalyzer with RNA 6000 Nano kit (Agilent Technologies). RNA concentration was measured also by Nanodrop ND-1000 (Thermo Scientific).

### Microarray hybridization

One-color microarray-based gene expression analysis was applied to analyse the RNA samples, on the Agilent-026652 Whole Human Genome Microarray 4×44K v2 platform (Agilent Technologies) according to the manufacturer’s protocol. To scan microarrays Agilent G49000 DA SureScan Microarray scanner was used and subsequently data were extracted by Agilent Feature Extraction.

### Differential expression analysis

Raw data obtained from microarray scanning were processed using Bioconductor software packages in R environment. Briefly, fluorescence intensities were background-subtracted, log_2_-transformed, and quantile-quantile normalized to get gene expression. Based on the results of hierarchical clustering and PCA on samples, one array (ID *Melanoma_5*) was excluded from the subsequent steps of analysis. Low-intensity probes (featuring a log_2_expression below 6.3 in more than one melanoma sample), were filtered out of the expression matrix as probes targeting unexpressed genes. Overall, 13,455 probes out of 34,127 (∼ 40%) were retained at the end of the filtering procedure and their log_2_expression values were tested for differential expression using rank product statistics. In particular, *n* = 5 melanoma biological replicates were compared against the single reference represented by the healthy biological mRNA pool of *n* = 4 non dysplastic naevi (RankProd v3.18.0 Bioconductor package, one-sample design) [45–49]. P-values were adjusted for multiple comparisons and all genes with a *q*-value (Benjamini-Hochberg False Discovery Rate, BH-FDR) < 0.05 were deemed as differentially expressed genes (DEGs) [50]. Finally, an additional cut off on fold changes (FCs) was applied to expunge from DEG lists genes having a |log_2_FC| < 0.5.

### Enrichment analysis

ToppFun web tool (by ToppGene Suite, https://toppgene.cchmc.org/) was used to analyse DEG lists for functional enrichment through a hypergeometric hypothesis test [51]. All terms with a BH-FDR *q*-value < 0.05 were considered statistically significant. Gene Set Enrichment Analysis was performed using GSEA v4.2.2 [51,52]. Expression data from microarray experiment were provided in the form of a pre-ranked list of genes (log_2_FC metric). Probes were collapsed to unique gene symbols before the analysis and a standard (weighted) enrichment statistic was chosen. Normalized Enriched Score (NES) and BH-FDR *q*-values are reported in the text for each gene set of interest. Within the context of GSEA, the threshold on *q*-value for a gene set to be considered statistically significant was set to 0.25. To evaluate the global transcriptional alterations affecting ribosomal proteins (rProteins), a custom gene set including all rProtein and rRNA genes was used. Details about such a custom Ribosomal Protein Gene Set (RPGS) have been already provided elsewhere [15].

### TCGA/GTEx validation

To provide an external validation of our main findings we used UCSC Xena Browser (University of California, Santa Cruz, http://xena.ucsc.edu/) [16] which allows the direct comparison of tumour expression data stored in TCGA with healthy samples from GTEx database (https://gtexportal.org/home/) [17]. Specifically, we filtered TCGA data in order to keep samples only from the skin cutaneous melanoma study (SKCM), of metastatic type (TM), excised from patients not harbouring any *BRAF* mutation. As for the control group, all the normal-skin samples retrieved from GTEx could be used. This led to a final comparison between *n* = 179 tumour samples and *n* = 557 normal tissues.

## Supplementary Materials

The following are available online at www.mdpi.com/xxx/s1, Figure SM1; Table SM1

## Author Contributions

Conceptualization, Elena Lastraioli and Annarosa Arcangeli; Data curation, Elena Lastraioli and Federico Alessandro Ruffinatti; Formal analysis, Elena Lastraioli and Federico Alessandro Ruffinatti; Funding acquisition, Annarosa Arcangeli, Francesco di Costanzo and Luca Munaron; Investigation, Elena Lastraioli and Federico Alessandro Ruffinatti; Methodology, Elena Lastraioli and Federico Alessandro Ruffinatti; Project administration, Elena Lastraioli and Federico Alessandro Ruffinatti; Resources, Francesco di Costanzo; Supervision, Annarosa Arcangeli, Francesco di Costanzo and Luca Munaron; Visualization, Elena Lastraioli and Federico Alessandro Ruffinatti; Writing – original draft, Elena Lastraioli and Federico Alessandro Ruffinatti; Writing – review & editing, Elena Lastraioli, Federico Alessandro Ruffinatti, Luca Munaron and Annarosa Arcangeli. All authors have read and agreed to the published version of the manuscript.

## Institutional Review Board Statement

the study was approved by the local Ethical Committee of Azienda Ospedaliero-Universitaria Careggi (BIO.16.028 released on October 5^th^ 2016)

## Informed Consent Statement

Informed consent was obtained from all subjects involved in the study.

## Data Availability Statement

Data are available upon reasonable request.

## Acknowledgments

The authors thank surgeons and nurses of the surgical compartment, Dr Luisa Di Cerbo, Prof Lorenzo Antonuzzo, Prof Lorenzo Leoncini, Prof Michele Maio and Dr Felice Arcuri for coordinating sample collection.

