## Supplementary figures and images for "Transcriptional landscape in *BRAF* wild type metastatic melanoma"

### Figure SM1

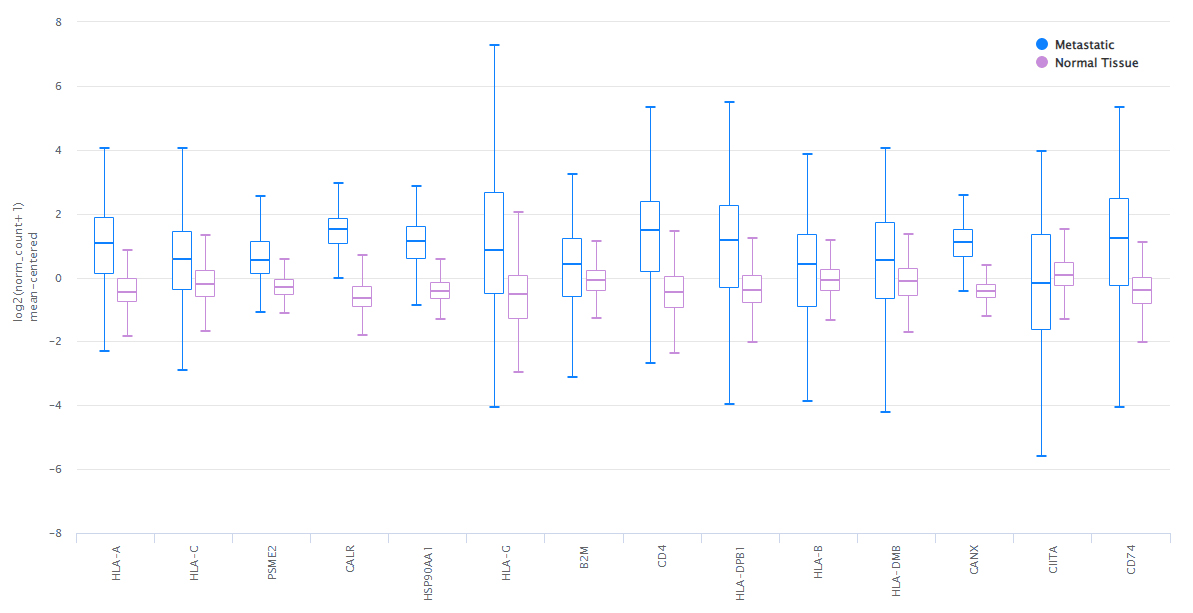
